# EGFR suppression and drug-induced potentiation are widespread features of oncogenic RTK fusions

**DOI:** 10.1101/2025.11.19.689362

**Authors:** Yuzhi (Carol) Gao, David Gonzalez-Martinez, Sofia Wissert, Hana Bader, Nidhi Sahni, Anh Le, Robert C. Doebele, Lukasz J. Bugaj

**Affiliations:** Department of Bioengineering, University of Pennsylvania, Philadelphia, PA 19104, USA; Department of Epigenetics and Molecular Carcinogenesis, Department of Bioinformatics and Computational Biology, The University of Texas MD Anderson Cancer Center, Houston, TX 77030, USA; Division of Medical Oncology, University of Colorado School of Medicine, Aurora, CO 80045, USA; Abramson Cancer Center, University of Pennsylvania, Philadelphia, PA 19104, USA; Quantitative and Computational Biosciences Program, Baylor College of Medicine, Houston TX 77030, USA; Institute of Regenerative Medicine, University of Pennsylvania, Philadelphia, PA 19104, USA

## Abstract

Regulation of cancer cells by their environment contributes to tumorigenesis and drug response, though the extent to which the oncogenic state can alter a cell’s perception of its environment is not clear. Prior studies found that EML4-ALK, a receptor tyrosine kinase (RTK) fusion oncoprotein, suppresses transmembrane receptor signaling through EGFR. Moreover, suppression was reversed with targeted ALK inhibition, thereby promoting survival and drug tolerance. Here we tested whether such modulation of EGFR was common among other RTK fusions, which collectively are found in ∼5% of all cancers. Using live- and fixed-cell microscopy in isogenic and patient-derived cell lines, we found that a wide variety of RTK fusions suppress transmembrane EGFR and sequester essential adaptor proteins in the cytoplasm, as evidenced by the localization of endogenous Grb2. Targeted therapies rapidly released Grb2 from sequestration and potentiated EGFR. Synthetic optogenetic analogs of RTK fusions confirmed that cytoplasmic sequestration of Grb2 was sufficient to suppress perception of extracellular EGF and could do so without driving signaling from the synthetic fusion itself, demonstrating that fusion signaling and suppression of EGFR could be functionally decoupled. Our study uncovers that a large number of RTK fusions simultaneously act as both activators and suppressors of signaling, the mechanisms of which could be exploited for new biomimetic therapies that enhance cell killing and suppress drug tolerance.

## Introduction

Interactions between cancer cells and their environment are central to disease progression and response to therapy and are therefore important to understand^1^. Recent studies have found that certain oncogenes can reshape a cell’s perception of environmental cues, in addition to their more established role of driving tonic oncogenic signaling^2–4^. Because such altered perception can contribute to pathology, it is critical to better understand the extent of oncogene-induced misperception and the mechanisms thereof.

Receptor tyrosine kinase (RTK) fusion oncoproteins are a large class of oncogenes found across ∼5% of human cancers^5–7^. Despite variable genetic composition, fusions share a common protein-level architecture: the intracellular domain of an RTK is fused to an oligomeric partner fragment from an unrelated protein. Whereas normal RTKs are embedded within the plasma membrane, fusions often lack the transmembrane domain associated with the RTK region, resulting in a multimeric chimera that localizes away from the plasma membrane. Nevertheless, cytoplasmic multimerization of the fusion is sufficient to activate the RTK fragment and subsequent tonic oncogenic signaling, most commonly through the Ras/Erk pathway.

EML4-ALK is an RTK fusion found primarily in non-small-cell lung cancer, accounting for ∼5% of cases. In addition to its ability to drive downstream signaling, EML4-ALK was recently found to suppress transmembrane receptor signaling^2^ (**Figure 1A,B**). Suppression occurred because of competitive inhibition: active EML4-ALK recruited adapters such as Grb2 that are required to transmit signaling both from the fusion as well as from transmembrane RTKs, including from EGFR. Consequently, EGFR was suppressed because active EML4-ALK sequestered adapters in the cytoplasm^2^, suppressing ∼90% of the information content from transmembrane receptor channels^8^. EML4-ALK is thus simultaneously both an activator of its own signaling and a suppressor of the signaling from the transmembrane compartment. Whether these dual suppressive and stimulatory activities can be decoupled is currently not known.

**Figure 1:**
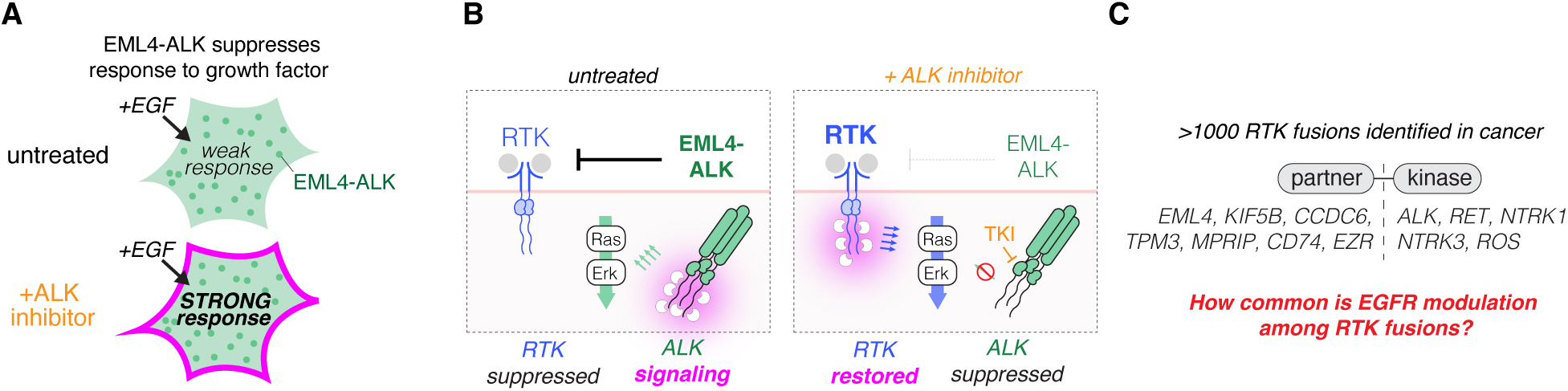
Modulation of EGFR signaling by EML4-ALK and potentially other RTK-fusion oncogenes. **A.** EML4-ALK fusion-driven cancer cells are insensitive to extracellular ligand stimulation. Targeted inhibition of oncogenic ALK signaling with tyrosine kinase inhibitors (TKIs) restores ligand sensitivity**. B.** EML4-ALK fusions form cytoplasmic assemblies that sequester key adaptor proteins, competitively inhibiting signaling by transmembrane receptor tyrosine kinases (RTKs). Pharmacological blockade of ALK activity releases these adaptors, resensitizing RTKs to ligand activation. **C.** Over 1,000 RTK fusion oncoproteins—comprising a kinase domain fused to a multimeric partner—have been implicated in cancer. The extent to which other fusions in this class sequester adaptors and suppress transmembrane RTKs is not clear.

The sequestration of adapters by EML4-ALK activity may have relevance for therapy. Treatment with tyrosine kinase inhibitors (TKIs) of ALK activity induce positive responses in patients with ALK rearrangements, although resistance to therapy often develops. In EML4-ALK+ cancer cells, ALK inhibition rapidly released sequestered adapters and potentiated transmembrane EGFR signaling, a well-established route of drug resistance^6,9–14^ (**Figure 1A,B**). Co-therapies that suppressed EGFR activity during ALK inhibition increased the efficacy of cell killing and suppressed therapeutic resistance, both in vitro and in mouse models^3,9,14–16^.

Because EML4-ALK shares a similar composition with other RTK fusions, the mechanisms of adapter sequestration and drug-mediated potentiation of EGFR may be shared across a substantial set of other RTK fusion-positive cancers, although such functional similarities have not been tested. Additionally, for EML4-ALK, the sequestration of adapters and suppression of EGFR co-occurred with the formation of large, micron-sized fusion condensates. However, it remains unclear whether such mesoscale condensates are required for sequestration Here we determine these functional principles of RTK fusion signaling using a panel of oncogenic RTK fusions, patient-derived cell lines, and engineered synthetic fusions (**Figure 1A-C**). We found that Grb2 sequestration, EGFR suppression, and drug-induced resensitization are widespread across fusions composed of a variety of RTK and partner fragments, although mesoscale condensation was rare. Finally, optogenetic fusions demonstrated that adapter sequestration can be decoupled from stimulation of downstream signaling and is sufficient to suppress EGFR.

## Results

### RTK fusions form assemblies with diverse subcellular distributions and suppress EGFR signaling

We explored functional commonalities among a panel of 10 distinct oncogenic RTK fusions, chosen to represent a variety of kinase and partner fragments and for their expression in available patient-derived cell lines (**Table S1**). Expression of GFP-tagged fusions in HeLa cells revealed an array of localization patterns (**Fig. 2A**). EML4-ALK Variants 1 (V1) and 3 (V3) showed cytoplasmic condensates, with V3 also displaying weak localization to microtubules, consistent with prior reports^17,18^. KIF5B-ALK and KIF5B-RET showed weak localization to microtubules and a strong polarization to peripheral structures. MPRIP-NTRK1 and EZR-ROS1 localized to actin-like structures, and CD74-ROS1 localized to the ER, as reported previously^19^. The remaining fusions (CCDC6-RET, TPM3-NTRK1, TPM3-ROS1) showed largely diffuse expression throughout the cytoplasm (**Fig. 2A**).

**Figure 2.**
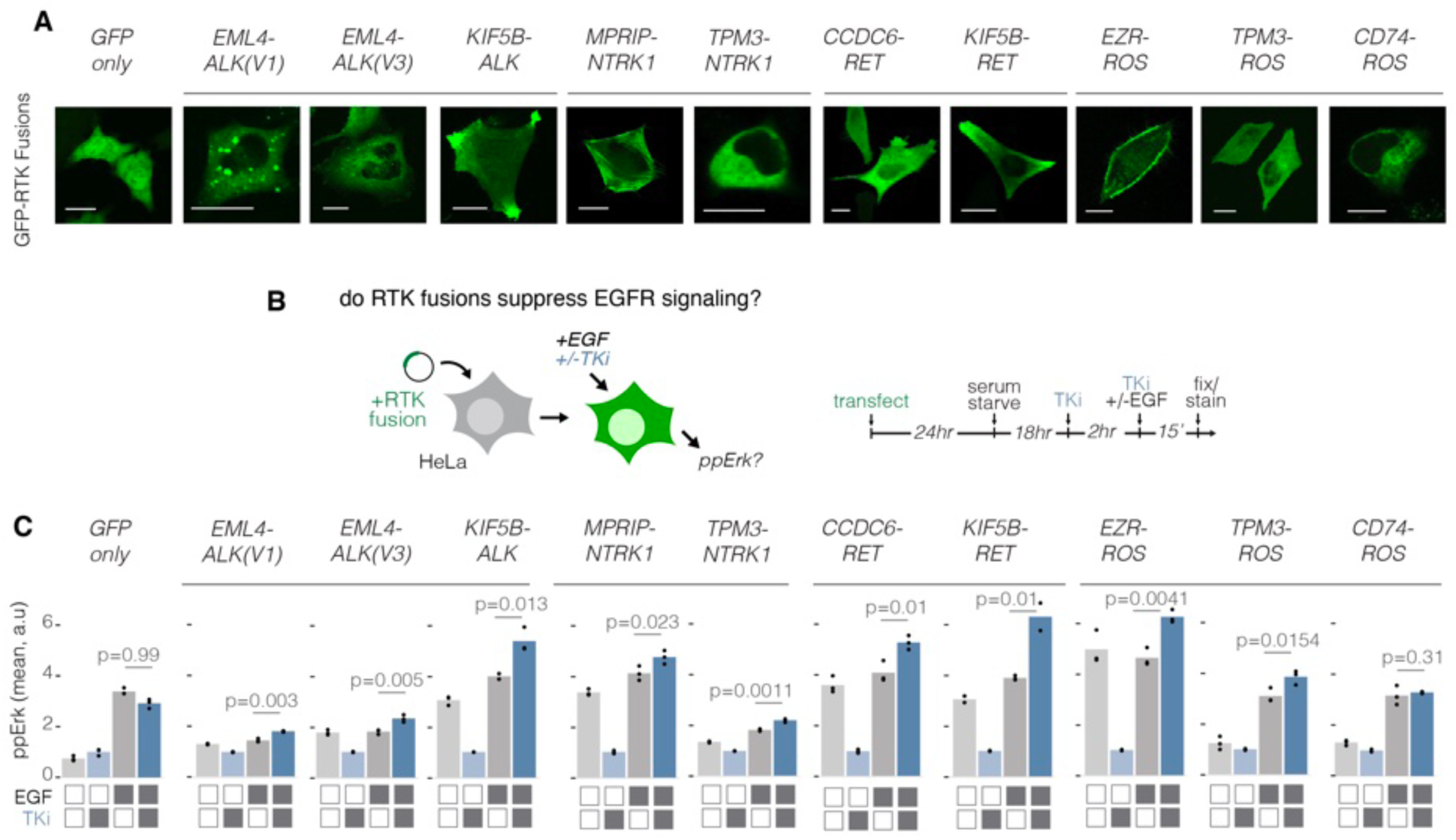
RTK fusions that signal through Ras-Erk exhibit variable subcellular localization but uniformly suppress EGFR signaling. **A.** Live-cell confocal images of HeLa cells expressing GFP-RTK fusions. Each panel shows a representative image from at least 20 analyzed cells per condition, with ≥3 independent experimental replicates. Scale bar = 20 µm. **B.** HeLa cells were transiently transfected with GFP-tagged RTK fusions or GFP-only control plasmids, treated with 1 µM TKI for 2 hours, followed by strong EGF stimulation (50 ng/mL, 15 min). **C.** Quantification of ppErk in RTK-fusion-expressing HeLa cells in response to EGF, with or without TKI pretreatment (100 nM–1 µM). Data are normalized to the mean of inhibited and unstimulated controls. Significance determined by one-sided t-test; n = 3 biological replicates, each representing 25-125 cells.

Next, we measured the ability of the fusions to signal through the Ras-Erk pathway and to suppress signaling through EGFR. We transfected HeLa cells with the GFP-tagged fusions and compared Erk activation (phospho-Erk or ppErk) under each of 4 conditions: 1) untreated, 2) treated with the appropriate tyrosine kinase inhibitor (TKI), 3) stimulated with EGF, and 4) stimulated with EGF after pretreatment with TKI (**Fig. 2B**). As expected, expression of RTK fusions led to an elevation in ppErk levels (condition 1), which decreased upon treatment with TKI (condition 2) (**Fig. 2C**). EGF stimulation in untreated cells resulted in only modest increases in signaling vs unstimulated cells (condition 3)(**Fig. 2C**). By contrast, pre-treatment with TKI (condition 4) resulted in a significant increase in EGF-stimulated ppErk. The lone exception was cells expressing CD74-ROS1, which showed no increase in response to EGF as a function of TKI pre-treatment, similar to the GFP control. These results indicate that, for most RTK fusions, kinase activity results in a strong suppression of transmembrane EGFR signaling, similar to our previous observations for EML4-ALK. Notably, EGFR suppression (and TKI-induced potentiation) was observed across several fusions that did not form condensates, suggesting that formation of mesoscale condensates is not essential for EGFR suppression (**Fig 2A**). Together, these results indicate that, although distinct fusions have unique localizations and signaling strengths, virtually all show a suppressive effect on signaling through TM RTK signaling.

### Oncogenic RTK fusions suppress EGFR signaling in patient-derived cancer cell lines

We next tested if EGFR suppression could be observed in patient-derived cell lines that harbored the oncogenes in our fusion panel^9,20–27^ (**Fig. 3A, Table S1**). We subjected the cancer lines to the same 4 conditions as for the HeLa cells and compared ppErk via immunostaining (**Fig. 3A**). As before, treatment with TKI suppressed ppErk levels in all cell lines, indicating elevated basal levels of oncogenic Ras-Erk signaling. While stimulation with EGF (50 ng/mL) increased ppErk levels in untreated cells, pretreatment with TKI consistently led to still higher stimulated levels of ppErk relative to cells that were not pretreated (**Fig. 3B,C**). Again, the lone exception was the cell line that expressed CD74-ROS1 (CUTO23). These results are consistent with prior reports that CD74-ROS1 does not signal strongly through the Ras-Erk pathway^19^.

**Figure 3:**
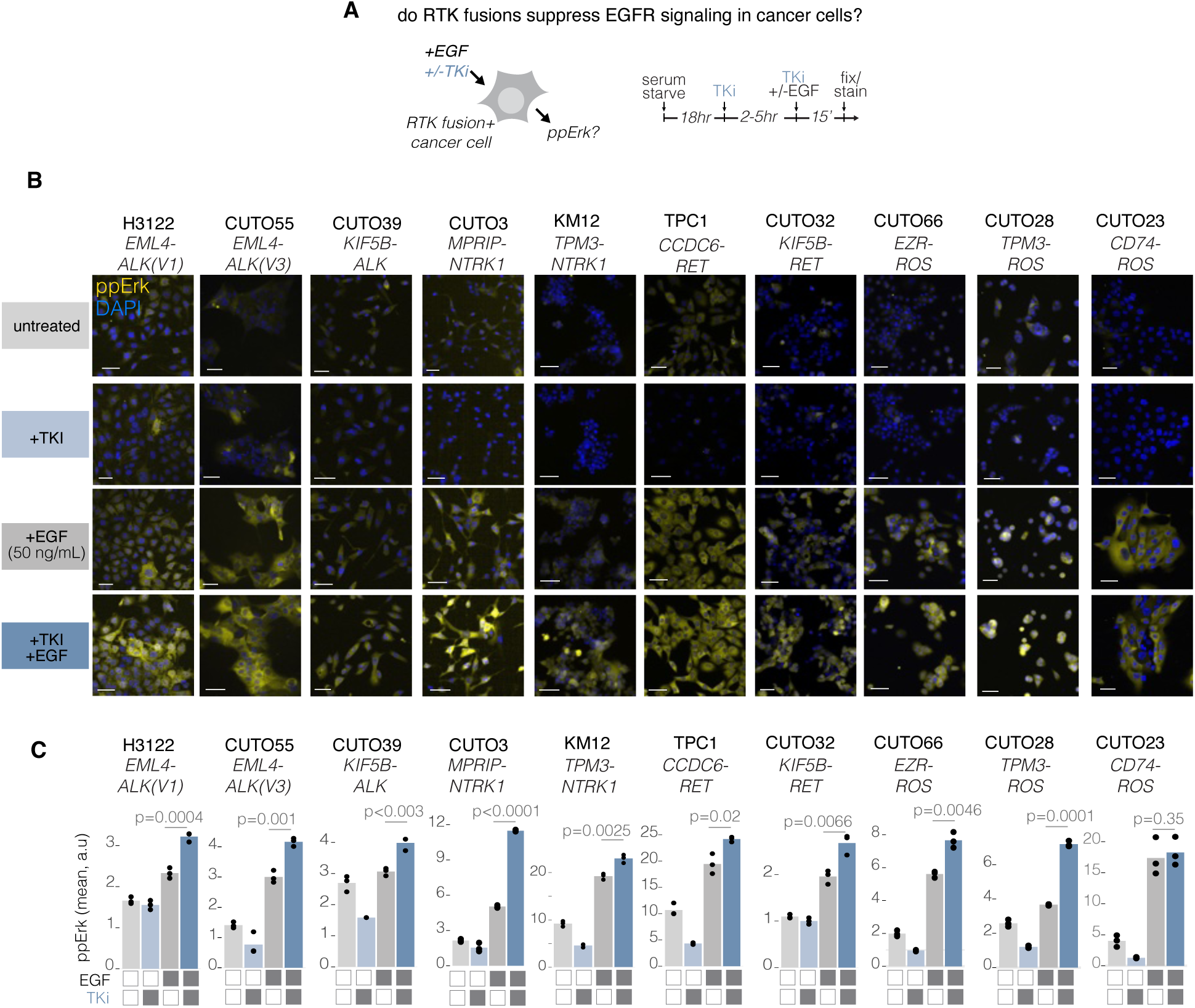
RTK-fusions suppress — and TKI treatment restores —EGFR signaling in patient-derived cancer cells. **A.** Cancer cells were pretreated with TKIs (100–1000 nM, 2–4 hours) before stimulation with EGF (50 ng/mL). **B.** Representative immunofluorescence images of ppErk in primary patient-derived cancer cell lines before and after EGF stimulation, with or without TKI pretreatment. Scale bar = 50 µm. **C.** Quantification of ppErk immunofluorescence in primary patient-derived cancer cell lines following EGF stimulation, with or without TKI pretreatment. All cell lines were pretreated with TKI for 2 hours prior to EGF stimulation, with the exception of TPC1 and CUTO39 cells, which were pretreated for 5 hours (**Fig. S2A-D**). Data points represent mean ppErk levels. Significance determined by one-sided t-test; n = 3 biological replicates, each representing 500-1000 cells.

Notably, of the three EML4-ALK V3 cell lines tested, two showed drug-induced sensitivity consistent with results from isogenic cells (**Fig 2C**), but one did not (CUTO34, **Fig S1**) suggesting additional factors in this cell line that can modulate EGF responsiveness in the presence of drug.

### RTK fusions sequester Grb2 and prevent its membrane translocation after EGF stimulation

Prior imaging and biochemical studies found that EML4-ALK suppresses EGFR signaling by binding downstream adapters like Grb2 and preventing their translocation to activated TM receptors, thereby suppressing transmission of TM signals^2^. We asked whether adapter sequestration could similarly explain the suppression of signal transmission in our panel of fusions (**Fig. 4A**). We expressed each mCherry-labelled fusion in lung epithelial Beas2B cells where endogenous Grb2 was tagged with mNeonGreen2 (mNG2), and we measured the extent of GRB2:mNG2 translocation to the membrane in response to EGF stimulation, the lack of which would indicate sequestration by RTK fusions (**Fig. 4B,C**). Strikingly, all fusions induced a near-complete suppression of Grb2 translocation in transfected vs untransfected cells, except for CD74-ROS1, which showed only mild suppression (**Fig. 4C-D, Movie S1-2**). For the fusions that efficiently sequestered Grb2, pretreatment with TKI reversed suppression, in most cases to levels observed in untransfected cells (**Fig. 4E**). A suppression score showed that TKI reduced fusion-associated suppression of EGFR by on average 75% (**Fig 4F,G**).

**Figure 4:**
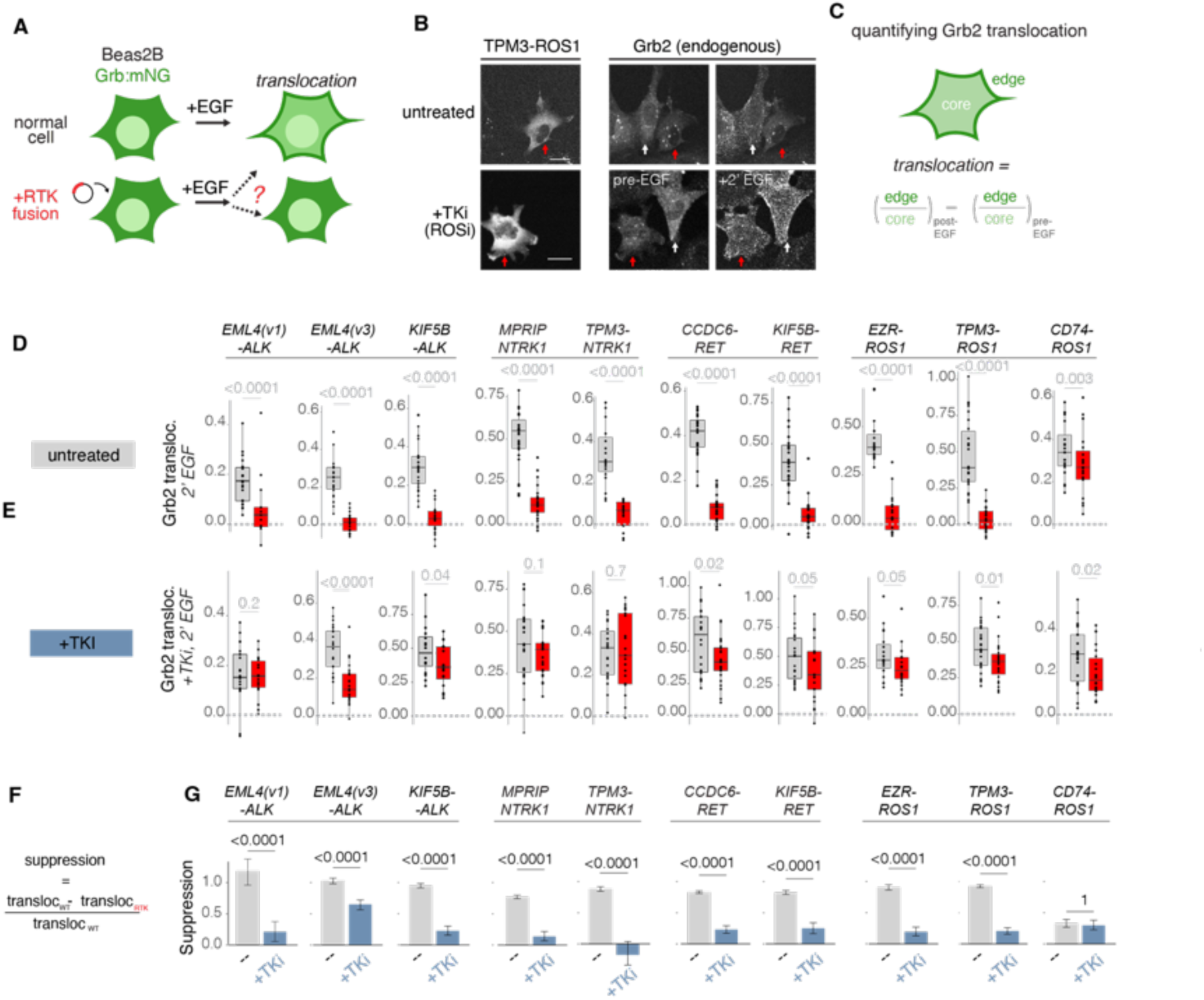
RTK fusions sequester GRB2 and prevent its translocation to the plasma membrane during EGF stimulation. **A.** GRB2:mNG Beas2B cells were transiently transfected with mCherry-tagged RTK fusions and stimulated with EGF (50 ng/mL) to assess Grb2 translocation in the presence or absence of an active fusion oncogene. **B.** Grb2 translocation is suppressed in cells expressing TPM3-ROS1 fusion but restored upon treatment with 1 µM entrectinib (ROSi) (**Movie S1**). White arrow indicates untransfected cells, and red arrow indicates transfected cells. Scale bar = 20 µm. **C.** Quantification method for Grb2 translocation. The cell edge was manually identified, and a 10-pixel cytoplasmic ring (“edge”) was defined. The remaining area was designated as the “core.” Membrane localization was calculated as the mean edge/core fluorescence ratio. Translocation was measured as the difference in membrane localization before vs. 2 min post-EGF stimulation. **D.** Grb2 translocation in cells transfected with mCherry-RTK fusions (red) vs. untransfected neighbors (grey). Each point represents a single cell. Boxplot elements: Median (line), quartiles (box), whiskers (1.5×IQR). **E.** Grb2 translocation in cells expressing mCherry-RTK fusions (cyan) vs. untransfected neighbors (grey) after 0.1-1 µM TKI treatment. **F.** Definition of translocation magnitude (difference in membrane localization pre-vs. post-stimulation). **G.** Comparison of Grb2 translocation suppression across RTK fusions. Data show median suppression from 1000 bootstrapped re-samplings. Error bars show lower and upper quartiles. Significance assessed by one-sided t-test.

In summary this series of experiments demonstrates that, independent of size, structure, or localization of their assemblies, RTK fusions largely share functional features, including the ability to signal through the Ras-Erk pathway, sequester RTK adaptors, and suppress transmembrane EGFR signaling. That CD74-ROS1 does not follow this trend may be explained by its localization to the ER, which exerts suppressive effects on RTK signaling^28,29^.

### Synthetic fusions show that Grb2 sequestration is sufficient to suppress EGFR and can be decoupled from fusion-driven signaling

Suppression of EGFR by cytoplasmic sequestration of adapters is a shared feature of RTK fusions, demonstrating the universal ability of fusions to simultaneously act as activators and suppressors of distinct sources of cellular signals. However, it was not clear whether adapter sequestration was sufficient to achieve EGFR suppression. It was also unknown whether EGFR suppression could be achieved without the concurrent stimulation of signaling downstream of the fusion, as observed for the naturally-occurring oncoproteins. To test these principles, we asked if we could engineer a minimal synthetic probe that sequestered Grb2 and suppressed EGFR but did not itself stimulate signaling. We thus sought to generate a minimal cytoplasmic RTK fragment that could bind and sequester Grb2 but itself might lack other necessary components for signaling. (**Fig. 5A-B**). We fused different RTK fragments to a variant of the Cry2 optogenetic clustering protein, Cry2_olig_^30,31^, which robustly forms visible protein clusters in cells (**Fig. 5C**). Although our studies showed that visible puncta are not necessary for EGFR suppression (**Figs. 2,3**), they nevertheless provide structures that allow visualization of adapter recruitment to activated proteins. We first tested cytoplasmic clustering of the intracellular domain (IC) of EGFR, since tyrosine phosphorylation (pY) of its tail generates Grb2 docking sites, yet EGFR itself does not commonly appear within cytoplasmic fusions. While light triggered condensation of Cry2_olig_-EGFR(IC) and subsequent recruitment of Grb2, it also triggered downstream Ras-Erk signaling, albeit only at intermediate expression levels (Fig. S2A-D, MovieS3-4).

**Figure 5:**
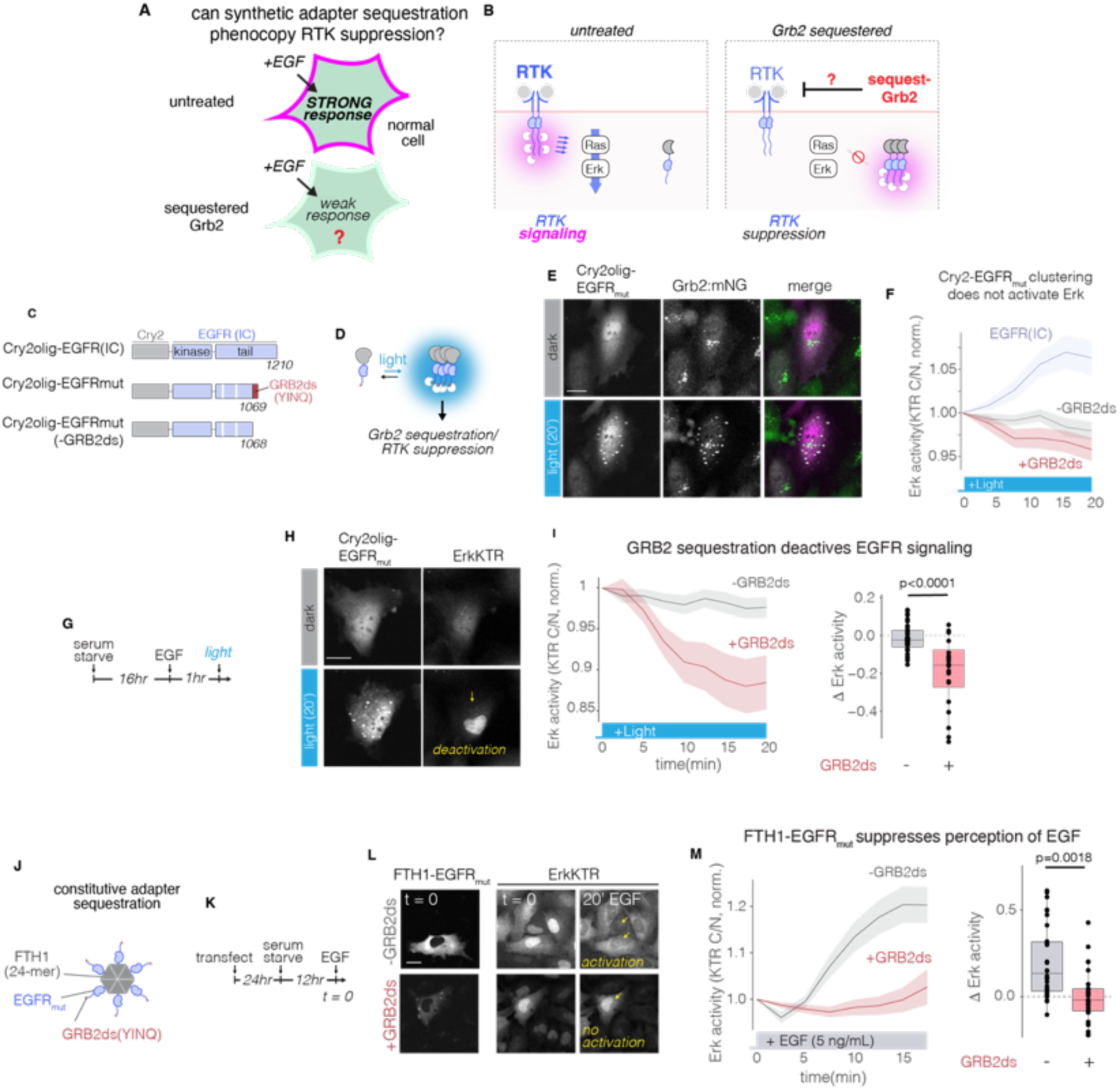
Adaptor sequestration is sufficient to modulate EGFR signaling. **A.** Determining whether Grb2 sequestration is sufficient for EGFR suppression. **B.** Inducing Grb2 sequestration by cluster-induced phosphorylation of Grb2 docking sites in the cytoplasm. **B.** Schematic of constructs to test optogenetic Grb2 sequestration. Cry2_olig_-EGFR: the entire intracellular domain (IC) of EGFR is fused to Cry2_olig_. Cry2_olig_-EGFR_mut_: same as Cry2_olig_-EGFR but the EGFR tail is truncated to the 1069 position (numbering of WT human EGFR). Additionally, Y998 and Y1016 are mutated to F (Fig S2). Finally, a YINQ motif, a known Grb2 docking site (GRB2ds), is appended to the C-term of this truncated construct to restore Grb2 binding. Cry2_olig_-EGFR_mut_(-GRB2ds): Same as Cry2_olig_-EGFR_mut_ but lacking the YINQ motif, thus serving as a negative control for Grb2 binding. **D.** Schematic of optogenetic control of synthetic fusion clustering and adapter binding. **E.** Representative images of Cry2_olig_-EGFR_mut_ (magenta) and GRB2:mNG (green) before and after 20 minutes of blue-light stimulation in Beas2B cells. Scale = 20 µm. (**Movie S5**). **F.** Quantification of Erk activity (Erk-KTR) cells expressing constructs shown in (C), at expression levels where Cry2olig-EGFR(IC) stimulates signaling (see **Figure S2**). Data show mean +/- SEM. Erk-KTR cytoplasmic/nuclear (C/N) ratio is normalized to T = 0.**G.** Experimental design to test suppression of EGFR signals. **H.** Cry2_olig_-EGFR _mut_ clusters with light stimulation and subsequently deactivates Ras-Erk signaling in Beas2B cells, indicated by nuclear localization of Erk-KTR. Scale = 20 µm. (**Movie S6**). **I.** Left: Quantification of **H.** Data show mean +/- SEM. Erk-KTR cytoplasmic/nuclear (C/N) ratio is normalized to timepoint 0. Right: Change in Erk-KTR C/N ratio after 20 min of light stimulation. Data points represent individual cells. Boxplot elements: Median (line), quartiles (box), whiskers (1.5×IQR). Significance assessed using one-sided t-test. **J.** The EGFR_mut_ tail is fused to FTH1 to generate constitutive adaptor sequestration (FTH1-EGFR _mut_). **K.** Experimental design to test responsiveness to EGF in the presence of FTH1-EGFR_mut_. **L.** Representative images of Beas2B cells expressing FTH1-EGFR _mut_ and Erk-KTR, both pre- and post-EGF stimulation. Arrows indicate cells transfected with FTH1-EGFR_mut_. Scale = 20 µm. (**Movie S7**). **M.** Left: Quantification of **L**. Data show mean +/- SEM. Erk-KTR cytoplasmic/nuclear (C/N) ratio is normalized to timepoint 0. EGF (5 ng/mL) is added immediately after imaging the initial frame. Right: change in Erk-KTR C/N ratio after 20 min EGF stimulation. Data points represent individual cells. Boxplot elements: Median (line), quartiles (box), whiskers (1.5×IQR). Significance assessed using one-sided t-test.

We then sought to eliminate signaling from Cry2_olig_-EGFR(IC) but retain Grb2 binding through 1) truncating the EGFR tail to the 1068 position to remove downstream pY sites previously reported to interact with Grb2^32–34;^ 2) mutating two additional pY sites (Y998F, Y1016F), and 3) appending a single known Grb2 binding motif^35^ (YINQ) to the C-terminus (**Fig. 5C-D, Fig. S3A**). Upon light activation, this modified fusion (Cry2_olig_-EGFR _mut_) formed visible puncta that colocalized with endogenous Grb2 (**Fig. 5E, MovieS5)**. Importantly, despite Grb2 recruitment, clustering of this construct did not activate downstream Erk at any level of expression (**Fig. 5F, S2D**).

Next, we asked if Grb2 recruitment by activated Cry2_olig_-EGFR_mut_ could suppress EGFR. We pre-stimulated cells with a high concentration of EGF (50 ng/mL) to ensure activation of EGFR/Erk signaling, which was confirmed by cytoplasmic localization (i.e. activation) of the Erk-KTR reporter^36^. Remarkably, light stimulation triggered a rapid return of the reporter to the nucleus despite the continued presence of EGF, indicating successful Grb2 sequestration and suppression of EGFR. By contrast, a control construct lacking the Grb2 docking site (-GRB2ds) showed no deactivation (**Fig. 5F-J, MovieS6**). Thus, Grb2 recruitment to Cry2_olig_-EGFR_mut_ is sufficient to outcompete EGFR for Grb2 and suppress its signaling, even in the presence of high concentrations of external ligands.

To further generalize our findings, we tested whether we could generate a constitutive, non-inducible construct that similarly suppressed EGFR. We replaced Cry2_olig_ with ferritin heavy chain (FTH1)^37^, a protein subunit that forms stable 24-mers (FTH1-EGFR _mut_). We first measured the dose-response to EGF of cells expressing FTH1-EGFR _mut_ or control constructs, including a fusion lacking the Grb2 site (-GRB2ds, **Fig. 5K**). FTH1-EGFR_mut_ suppressed sensitivity to EGF across all concentrations relative to controls (**Fig. S3B-D**). We validated this result using live-cell confocal microscopy, finding suppressed EGF-induced Erk activity in cells expressing FTH1-EGFR _mut_ compared to the - GRB2ds control (**Fig. 5L-O, MovieS7**).

Collectively, we find that sequestration of Grb2 in the cytoplasm is sufficient to achieve transmembrane suppression and can be achieved without concurrent stimulation of signaling from the cytoplasm.

## Discussion

Our study uncovers fundamental principles of RTK fusion signaling through an extensive characterization of a variety of molecularly distinct fusions as well as through construction of minimal synthetic fusions. We confirmed known shared features of fusions, such as the ability to stimulate downstream Ras-Erk signaling. We also found additional commonalities, including the ability of signaling-competent fusions to sequester Grb2 and to suppress signaling through transmembrane EGFR, a behavior previously described only for EML4-ALK. The presence of EGFR suppression among diverse active fusions —coupled with its absence among active transmembrane receptors^2^—suggests that suppression is a specific property of RTKs that are not anchored to the plasma membrane.

Prior studies suggested that assembly of RTKs and downstream adapters into condensates may be a causal mechanism for signal transmission from cytoplasmic fusions to downstream pathways^18,38–40^. However, mesoscale condensation was absent from most members in our panel, which displayed variable distribution patterns including localization to subcellular compartments, polarization, and diffuse distribution in the cytoplasm, indicating that condensation is not necessary for either signaling or EGFR suppression. Our results from the synthetic EGFR_mut_ fusion, which recruited Grb2 into condensates but did not trigger signaling, further indicate that assembly of Grb2 into condensates of active kinase is not sufficient for signaling. Further studies are needed to assess what additional factors are required to produce downstream signals and to understand the specific role of condensates in signal transmission. The ability of Cry2-EGFR(IC)—but not Cry2-EGFR_mut_— to trigger signaling provides a testbed for future studies to determine how cytoplasmic RTK fusions transmit oncogenic signals.

The ability of fusions to suppress EGFR but potentiate it during drug treatment carries therapeutic implications. In the case of EML4-ALK-driven cancer cells, the relief of sequestration after ALK inhibition led to rapid reactivation of Ras-Erk signaling and subsequent drug tolerance, and tolerance was suppressed by co-treatment with inhibitors that blocked EGFR reactivation^2,9^. Because we find the same underlying adapter and signaling dynamics across all fusions that signal through Ras-Erk, cancers driven by these fusions may all benefit from co-therapies that suppress rapid EGFR reactivation. Furthermore, our ability to isolate the EGFR-suppressive nature of fusions and recapitulate it in minimal synthetic constructs suggests that targeted sequestration of signaling adapters could be harnessed as a novel strategy to broadly suppress transmembrane RTK reactivation, which could find use against a range of cancers where receptor hyperactivation drives disease or limits therapeutic response.

## Acknowledgements

This work was supported by the National Institutes of Health (R35 GM138211 for L.J.B. and D.G.M.), the American Cancer Society (RSG-22-176-01-TBE to L.J.B.), and CPRIT (RP220292 to N.S.). Cell sorting was performed on a BD FACSAria Fusion that was obtained through NIH S10 1S10OD026986.

## Author Contributions

Y.C.G., and D.G.M., and L.J.B. conceived the study. Y.C.G. performed experiments and D.G.M provided guidance on imaging and analysis techniques. N.S. bioinformatically prioritized the RTK fusions used for experiments and provided input on the study design. Y.C.G., H.B., and S.W. cloned constructs. A.L. and R.C.D. derived the CUTO cell lines from patient tumors. Y.C.G analyzed data. L.J.B. provided experimental feedback and guidance. Y.C.G. performed statistical analysis. L.J.B. supervised the work. Y.C.G. and L.J.B. wrote the manuscript and made figures, with editing from all authors.

## Methods

### Cell lines and cell culture

All cell lines were maintained at 37°C with 5% CO₂ in a humidified incubator. Patient-derived thyroid cancer lines (TPC-1, KM12, CUTO-23/28/32/34/39/46/55/66) were cultured in RPMI-1640 (Gibco #11875093) supplemented with 10% FBS (Gibco #10437028) and 1% penicillin/streptomycin (Gibco #15140122). HeLa cells were grown in DMEM (Gibco #11965092) with identical supplements. Beas2B GRB2:mNG cells (generated via CRISPR-Cas9 knock-in of mNeonGreen at the endogenous *GRB2*locus) were cultured as above. The CUTO series were generous gifts from Dr. Robert Doebele (University of Colorado, Denver, CO). The generation of Beas2B ErkKTR-iRFP/H2B-BFP cells was achieved by transfection with a PiggyBac transposon system^41^, composed of transfer plasmids containing the inserts and a Super piggyBac Transposase expression vector (System Bio-sciences, PB210PA-1), followed by subsequent puromycin selection and fluorescence-activated cell sorting (FACS).

For experiments, cells were seeded in fibronectin-coated (10 μg/mL, MilliporeSigma #FC010) 384-well plates (Corning #354663) and, unless indicated otherwise, cells were serum starved overnight before experiments by performing 8x washes with serum-free medium, either manually or using the BioTek 405 LS microplate washer.

### Reagents and inhibitors

All cell lines were pretreated with the following inhibitors for 2 hours prior to EGF stimulation, with the exception of TPC1 and CUTO39 cells, which were pretreated for 5 hours: crizotinib (1 μM, Sigma-Aldrich, PZ0191), alectinib (0.1 μM, Selleckchem, CH5424802), BLU-667(0.1 μM, ABCam, B2548), entrectinib(0.1 μM, Selleckchem, S7998). Cells were stimulated with EGF (50 ng/ml unless stated otherwise, PeproTech, 315-09).

### Plasmid design and assembly

All cloning was performed by PCR and DNA assembly using NEBuilder® HiFi DNA Assembly Master Mix (New England Biolabs #E2621) or KLD Enzyme Mix reaction (New England Biolabs #M0554). EML4-ALK(V1) and EML4-ALK(V3) were obtained from previously described constructs. The CCDC6-RET construct was a generous gift from the Dr. Richard Kriwacki (St. Jude Research Hospital). The 5 RTK Gene fusions KIF5B-ALK, KIF5B-RET, TPM3-NTRK1, CD74-ROS1 and TPM3-ROS1 were identified by computational (TopHat-Fusion) analysis of transcriptome sequencing data from the Cancer Genome Atlas (TCGA) (RRID:SCR_003193). To obtain fusion proteins, we filtered the list for “in-frame” gene fusions (*i.e.*, chimeric proteins made from joining two genes that usually encode separate proteins). Their breakpoints or junctions were prioritized based on the top frequencies of occurrence found in TCGA: KIF5B-ALK: chr10:32022140 (KIF5B) and chr2:29223528 (ALK); KIF5B-RET: chr10:32028428 (KIF5B) and chr10:43116584 (RET); TPM3-NTRK1: chr1:154170400 (TPM3) and chr1:156874571 (NTRK1); CD74-ROS1: chr5:150404680 (CD74) and chr6:117324415 (ROS1); TPM3-ROS1: chr1:154170400 (TPM3) and chr6:117321394 (ROS1).KIF5B-ALK, KIF5B-RET, TPM3-NTRK1, CD74-ROS1, and TPM3-ROS1, and EZR-ROS1 plasmids were cloned into pCMV-mGFP plasmid. The KIF5B-ALK plasmid was then modified using PCR followed by KLD reaction (New England Biolabs #M0554S) to match the variant (K17;A20) found in the CUTO39 cancer cell line. The MPRIP-NTRK1 plasmid was constructed by fusing multiple DNA fragments synthesized by Twist Biosciences using HiFi DNA Assembly Master Mix based on the previously published sequence^26^. All RTK fusions were also cloned downstream of mCherry in a pCMV-mCh plasmid. For live-cell tracking of ERK activity, we generated stable reporter cell lines using the PiggyBac transposon system (System Biosciences). pBRPB-CAG-ErkTKR-iRFP was generated by inserting ErkKTR-IRFP coding sequence in place of mCherry in pBRPB - CAG-mCherry-IP (Addgene:#106333). Visualization of nuclei was achieved using a pBRPB-CAG-H2B-BFP plasmid. For FTH1-EGFR and Cry2_olig_-EGFR constructs, EGFR variants were generated using site-directed-mutagenesis and inserted downstream of pHR-mCh-FTH1 or pHR-mCh-Cry2_olig_.

### Transient transfection

Beas2Bs and HeLa cells are transfected using Fugene4K (Promega, E5911). according to the manufacturer’s protocol. Briefly, 2500 to 5000 cells were seeded in fibronectin-coated wells of a 384-well plate in RPMI or DMEM growth medium. Each well as supplemented with 25ng of DNA plasmid and 0.75uL of the Fugene4K transfection reagent, diluted in OptiMEM (Gibco, #31985070).

### Optogenetic stimulation

Light stimulation was performed on a Nikon Ti2-E microscope 488nm laser, with 300ms exposure and stimulated every 30 seconds.

### Growth factor stimulation

Unless otherwise indicated, starved cells were treated with the indicated inhibitors for 2 hours and then stimulated at their respective time points with EGF. Cells were then fixed simultaneously for 10 minutes with 4% paraformaldehyde (PFA) (Electron Microscopy Sciences, 15710). Samples were then processed for immunostaining.

### Immunostaining

Fixed cells were permeabilized with 0.5% Triton-X100 in PBS for 10 min, followed by incubation in ice-cold 100% methanol for 10 min. Samples were then blocked with blocking solution (1% bovine serum albumin (BSA, Fisher, BP9706100) diluted in PBS) for 1 hour at room temperature. Samples were incubated in indicated primary antibody diluted in blocking solution for either 2 hours at room temperature (RT) or overnight at 4 °C. Primary antibodies used were: phospho-p44/42 MAPK (Erk1/2) (Thr202/Tyr204), Cell Signaling #4370. Antibodies were used at dilutions recommended by the manufacturer. After incubation with primary antibody, samples were washed 5X with 80% washes of 0.1% Tween-20 in PBS (PBS-T). Samples were then incubated in blocking solution containing secondary antibody (IgG (H+L) Cross-Adsorbed Goat anti-Rabbit, DyLightTM 488, Invitrogen #35553; Goat anti-Rabbit IgG (H+L) Cross-Adsorbed Secondary Antibody, DyLightTM 650, Invitrogen #SA510034; and 4,6-diamidino-2-phenylindole (DAPI; ThermoFisher Scientific #D1306, 300 nM) for 1 hour at RT. Samples washed with PBS-T as previously described.

### Live cell imaging

Live cell imaging was performed using a Nikon Ti2-E microscope equipped with a Yokagawa CSU-W1 spinning disk, 405/488/561/640 nm laser lines, an sCMOS camera (Photometrics), and a motorized stage. Cells were maintained at 37 °C and 5% CO2 using an environmental chamber (Okolabs). All imaged wells were cultured in phenol free RPMI. For adaptor localization assay in Beas2B GRB2:mNG2 cells, cells were serum-starved before imaging. For imaging of adaptor translocation in response to EGF stimulation, cells were imaged every 30 sec using a 40X oil-immersion objective.

### Image processing and analysis

Images of fixed-cell immunostaining for cancer cells were quantified using CellProfiler (v 4.0.7)^42^. Briefly, cell nuclei were segmented using the DAPI channel, and the cytoplasmic fluorescence was measured within a 5-pixel ring that circumscribed the nucleus. Images for fixed-cell immunostaining for isogenic cells overexpression oncogenes is segmented and whole-cell pErk intensity is quantified using CellPose^43^. Quantification was exported to R for processing and data visualization using the tidyR package^44^.

### GRB2 translocation

Time-lapse images of GRB2 translocation in response to EGF stimulation were quantified with a custom MATLAB script previously described. Briefly, Translocation was quantified by identifying the cell edge and defining a 10 pixel ring into the cytoplasm (“edge”). The remaining cell pixels beyond this ring were designated as the cell “core”. Membrane localization was defined as the ratio of mean edge fluorescence to mean core fluorescence. Translocation was defined as the difference in adapter membrane localization after 2 min post EGF stimulation vs pre-stimulation.

### Erk-KTR activity

Erk-KTR dynamics were tracked and quantitated in an automated manner using a customized pipeline Python script. Briefly, the H2B-BFP channel is used to segment nuclei with customized CellPose models. Next the nuclei segmentations are tracked between timepoints by calculating the amount of pixel overlap between objects in the current frame and matches in the previous frame. Then, to calculate cytoplasmic intensity of the ErkKTR and expressed constructs, a 5-pixel ring is created surrounding the segmented nuclei. Then the ErkTKR cytoplasmic/nuclear ratios from cells are calculated and imported into R Studio for visualization.

### Statistics

Means and p-values between two groups were compared using one-tailed t-tests. Medians of index of suppression were compared by a bootstrap test. From each original data set, 1000 new datasets of the same size were generated by resampling with replacement. The means of these resampled data were used to calculate the suppression metric. Thus 1000 suppression metrics were generated for each condition, allowing statistical comparison. Analyses were performed using R, version 4.0.3.^44^

---

**Table S1.** Patient derived cancer cell lines, associated RTK fusions, and TKIs conditions used in experiments.

| FUSION | CELL LINE | TKI USED |
| --- | --- | --- |
| **MPRIP-NTRK1(M21;N14)** | CUTO3 | entrectinib (100nM) |
| **CD74-ROS1 (C6;R34)** | CUTO23 | entrectinib (100nM) |
| **TPM3-ROS1 (T7;R35)** | CUTO28 | entrectinib (100nM) |
| **KIF5B-RET (K15;R12)** | CUTO32 | BLU-667 (100nM) |
| **EML4-ALK (E6;A20)** | CUTO34 | crizotinib (1µM) |
| **KIF5B-ALK (K17;A20)** | CUTO39 | crizotinib(1µM) |
| **EML4-ALK (E6;A20)** | CUTO46 | crizotinib(1µM) |
| **EML4-ALK (E6;A20)** | CUTO55 | crizotinib(1µM) |
| **EZR-ROS1 (E9;R34)** | CUTO66 | entrectinib (100nM) |
| **EML4-ALK (E13;A20)** | H3122 | crizotinib(1µM) |
| **TPM3-NTRK1(T7;N12)** | KM12 | entrectinib (100nM) |
| **CCDC6-RET(C3;R12)** | TPC-1 | BLU-667 (100nM) |

**Figure S1.**
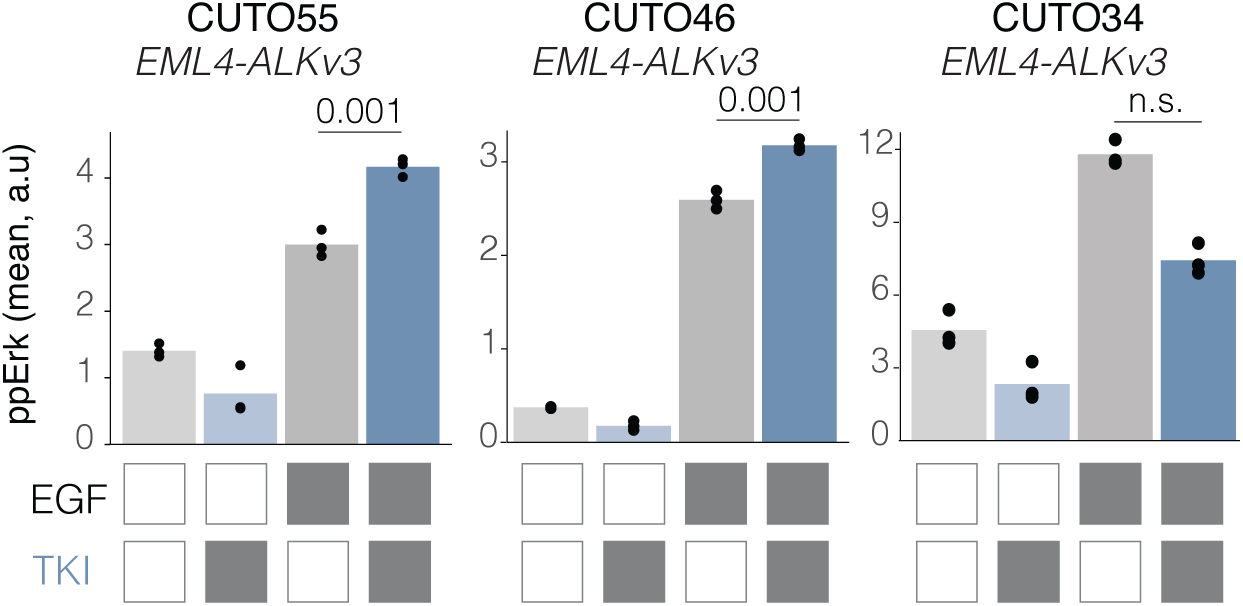
Comparison of EGFR resensitization among distinct patient-derived cell lines driven by EML4-ALK V3. Quantification of ppERK immunofluorescence in EML4-ALKv3+ cancer cell lines following EGF stimulation, with or without 2hr of TKI pretreatment. Data points represent mean ppERK levels. Significance determined by one-sided t-test; n = 3 biological replicates. CUTO55 data is identical to data from Fig.3E, included here for ease of comparison.

**Figure S2:**
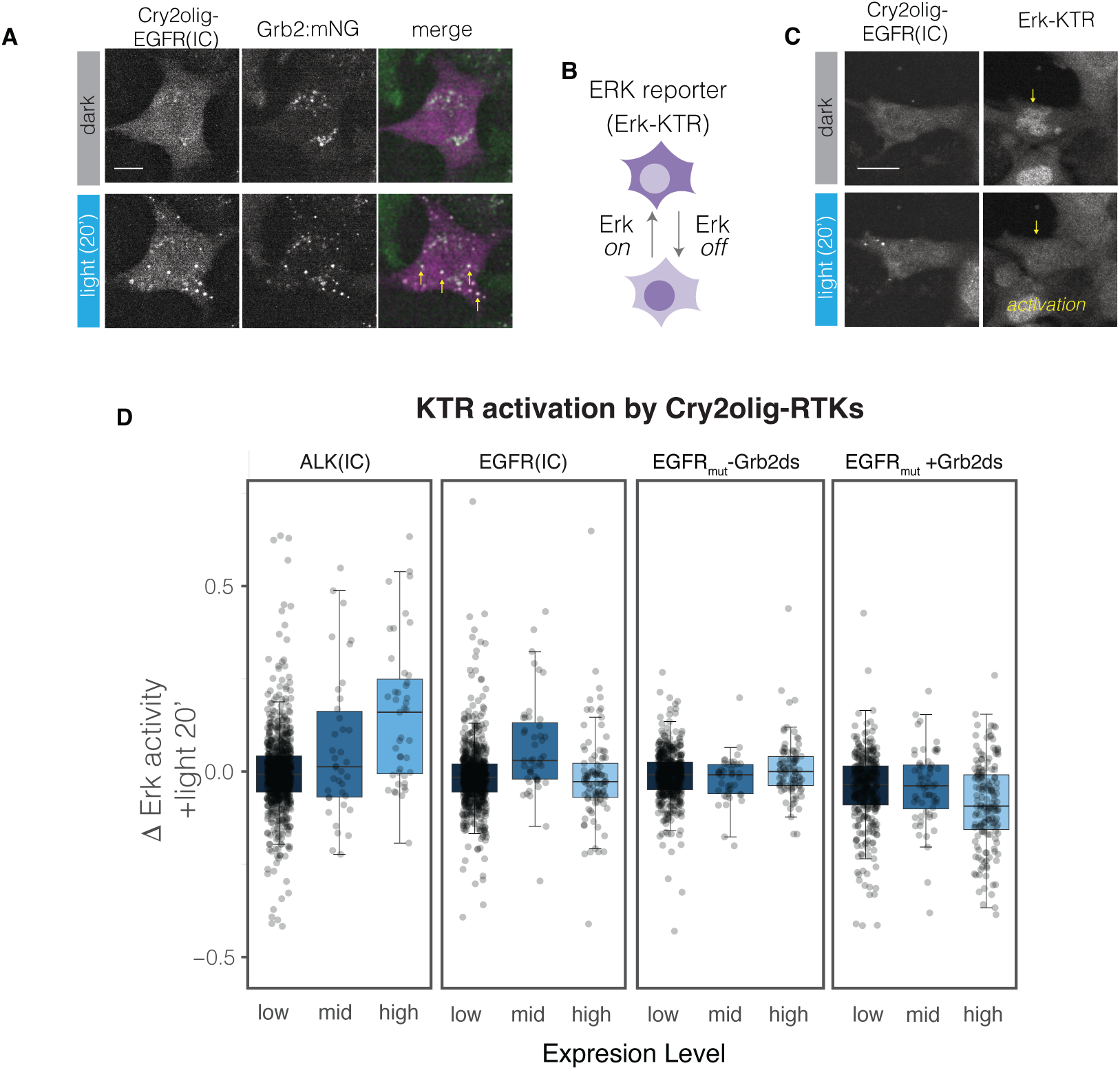
Multimerizing EGFR(IC) activates Ras-Erk signaling at intermediate expression levels, but multimerized EGFR_mut_ does not. **A.** Representative images showing localization of Cry2_olig_-EGFR(IC) (magenta) and GRB2:mNG (green) in Beas2B cells before and after 20 min blue light stimulation. Scale = 20 µm. See **Movie S3**. **B.** The Erk-KTR biosensor indicates ERK activity through nuclear-cytoplasmic translocation. **C.** Representative image showing Erk activation after light stimulation of Cry2_olig_-EGFR(IC) in Beas2B cells in serum-starved conditions. Scale = 20 µm. See **Movie S4**. **D.** Quantification of changes in Erk activity after 20 min of light stimulation of cells expressing Cry2_olig_-ALK(IC), Cry2_olig_-EGFR(IC), Cry2_olig_-EGFR_mut_ (-Grb2ds), and Cry2_olig_-EGFR_mut_ (+Grb2ds). Data points represent individual cells. Boxplot indicates the median and upper/lower quartiles, and whiskers extend to 1.5*IQR.

**Figure S3:**
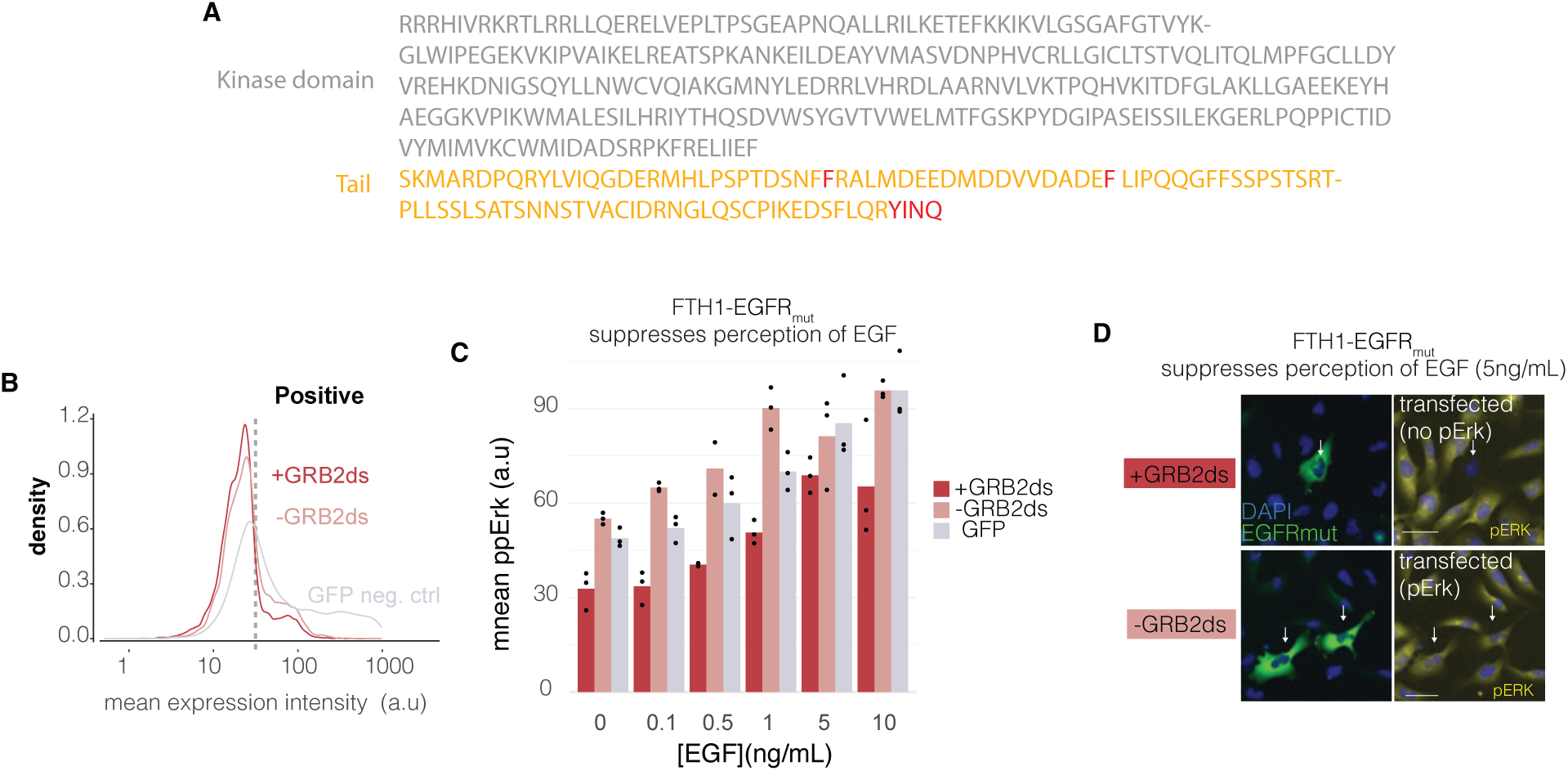
Multimerizing EGFR_mut_ suppresses EGFR signaling through the Ras-Erk pathway. **A.** Amino acid sequence of the EGFR_mut_ region. Grey indicates the kinase domain. Orange indicates the mutated EGFR tail. Residues marked in red are Y>F mutations and the appended YINQ motif (GRB2ds). **B.** Distribution of GFP-FTH1-EGFR_mut_(+/- GRB2ds) expression in Beas2B cells. Transfected cells are defined as having expression higher than threshold (dashed grey line). **C.** Quantification of ppERK in Beas2B cells expressing FTH1-EGFR _mut_ in response to a range of EGF dosages. Data points represent means of 50-300 cells per condition. **D**. Representative images of ppERK immunofluorescence (yellow) in Beas2B transiently expressing GFP-FTH1-EGFR_mut_(+/- GRB2ds) (green).

**Movie S1.** TPM3-ROS1 expression suppresses GRB2 translocation to the plasma membrane upon EGF ligand stimulation (50 ng/mL). **Time, mm:ss. Scale bar, 10 μm.**

**Movie S2. ROSi pretreatment restored GRB2 translocation to the plasma membrane in TPM3-ROS1-expressing cells upon EGF stimulation (50 ng/mL).** Time, mm:ss. Scale bar, 10 μm.

**Movie S3. Cry2_olig_-EGFR(IC) multimerizes and colocalizes with GRB2:mNG upon light stimulation.** Time, mm:ss. Blue box indicates light stimulation. Scale bar, 10 μm.

**Movie S4. Cry2_olig_-EGFR(IC) multimerizes and activates ERK signaling upon light stimulation.** Time, mm:ss. Blue box indicates light stimulation. Scale bar, 10 μm.

**Movie S5. Cry2_olig_-EGFR_mut_ multimerizes and colocalize with GRB2:mNG upon light stimulation.** Time, mm:ss. Blue box indicates light stimulation. Scale bar, 10 μm.

**Movie S6**. **Cry2_olig_-EGFR_mut_ multimerizes and deactivates pre-activated EGFR upon light stimulation.** Time, mm:ss. Blue box indicates light stimulation. Scale bar, 10 μm.

**Movie S7. FTH1-EGFR_mut_ suppresses ERK signaling responses to EGF stimulation. (5ng/mL).** Time, mm:ss. Scale bar, 10 μm.

